# Identification and characterization of UBA5 inhibitors

**DOI:** 10.1101/2024.10.25.620379

**Authors:** Vigyasa Singh, Yuansong Wan, Tahsin Reaz Ahmed, Si-Yu Wang, Subodh Kumar Samrat, Zhong Li, Subhash Sinha, Li Gan, Hongmin Li

**Author notes:** Corresponding author: Hongmin Li.

## Abstract

UBA5 is a critical E1-activating enzyme in the UFMylation pathway, a post-translational modification process implicated in neurodegenerative diseases and cancers. In this study, we developed a high-throughput screening (HTS) assay to identify inhibitors of UBA5 from a library of blood-brain barrier (BBB)-permeable compounds. The assay, based on the AMP-Glo™ kit, enabled the identification of five novel UBA5 inhibitors belonging to three distinct chemical scaffolds with low micromolar *IC_50_* values. These inhibitors demonstrated selectivity for UBA5 over other E1 enzymes, such as UBA1, and showed efficacy in inhibiting endogenous UFMylation in HEK293T cells. The identified inhibitors not only provided valuable tools for studying UFMylation but also represent potential therapeutic candidates for diseases associated with dysregulated UFMylation, such as Alzheimer’s disease and cancer. Further optimization of these compounds is necessary to enhance their potency and pharmacokinetic properties. Our findings highlight UBA5 as a promising therapeutic target and pave the way for the development of selective inhibitors that can modulate UFMylation in human diseases.

## INTRODUCTION

The ubiquitin-fold modifier 1 (UFM1) conjugation system, known as UFMylation, is an essential post-translational modification pathway that regulates a broad array of cellular processes, including protein folding, stress response, and DNA damage repair^1–6^. UFMylation is catalyzed by three core enzymes: the E1-activating enzyme UBA5, the E2-conjugating enzyme UFC1, and the E3-ligase UFL1, which work sequentially to attach UFM1 to target substrates. Disruption of this pathway has been linked to several diseases, including cancer^4,7–11^, neurodegenerative disorders^12–16^, and developmental diseases^17–19^, emphasizing the critical need to understand and potentially target this system therapeutically. Among these enzymes, UBA5 plays a pivotal role in initiating the UFMylation process by activating UFM1 through ATP-dependent adenylation and subsequent transfer to UFC1.

UBA5, as an E1-activating enzyme, occupies a unique position in the UFMylation cascade by catalyzing the first step of UFM1 activation. This function is crucial because it represents the commitment step toward UFMylation, controlling the overall flux of this modification in cells. Due to its central role, any perturbation in UBA5 function can have profound consequences on cellular physiology. For instance, UBA5 alterations have been implicated in a range of human diseases, including Alzheimer’s disease^12,13^ and various cancers^4,7^, where abnormal UFMylation either contributes to the pathogenesis or progression of these conditions. Hence, UBA5 has emerged as a potential therapeutic target in drug discovery efforts aimed at modulating the UFMylation pathway.

In recent years, evidence has accumulated pointing to UBA5’s involvement in neurodegenerative diseases, particularly Alzheimer’s disease^12–15^. Altered UFMylation dynamics, often due to UBA5 mutations and dysregulation, have been linked to impaired protein homeostasis, which is a hallmark of Alzheimer’s disease^20–23^. For instance, studies have identified mutations in the *UBA5* gene in patients suffering from early-onset neurodegeneration^24^, further supporting the idea that UFMylation contributes to the development of neurological disorders. Moreover, UBA5’s role in cellular stress response pathways, including endoplasmic reticulum (ER) stress^1–3,25–30^, highlights its importance in the maintenance of neuronal function under stress conditions, which is another critical factor in Alzheimer’s pathology^20–23^.

UBA5 has also been implicated in the pathogenesis of cancer, with studies showing that abnormal UFMylation can influence oncogenic signaling pathways. For example, dysregulated UFMylation has been linked to the development and progression of various cancers, including breast, colon, gastric, glioblastoma, liver, lung, oral, pancreatic, and renal cancers^4,11,31–36^. UBA5 overexpression has been observed in tumor samples from patients with these cancers, correlating with poor prognosis^4,7,11^ and increased metastatic potential^11,31^. The oncogenic role of UBA5 is thought to stem from its influence on key cellular processes, such as apoptosis, autophagy, and DNA repair^1–3,25–30^, all of which are often altered in cancer cells. Thus, UBA5 represents a promising target for the development of novel anti-cancer therapies.

Despite the significant role of UBA5 in human disease, only a limited number of inhibitors have been identified^11,37–39^. One identified inhibitor, adenosine 5’-sulfamate (ADS), is an ATP-competitive analogue with an *IC_50_* of 13 µM^37^. However, ADS is a pan-E1 inhibitor, limiting its utility^37^. Another inhibitor, named compound **8.5**, was developed as a selective UBA5 inhibitor with an *IC_50_*of 4 µM^38^. Despite its selectivity, compound **8.5** is a Zn^2+^-chelator, which may reduce its therapeutic potential. A third compound, DKM 2-93 (DKM), was discovered as a covalent UBA5 inhibitor, but it has a relatively high *IC_50_* of 430 µM^39^. The natural product Usenamine A has been found to influence UBA5 expression^11^; however, it remains unconfirmed whether UBA5 is its direct target. Currently, there is no evidence of a direct interaction between Usenamine A and UBA5, nor of any inhibition of UBA5 enzymatic activity by the compound. To date, no high-throughput screening (HTS) efforts have been reported for UBA5 inhibitors.

In this manuscript, we report the development of an HTS assay to identify UBA5 inhibitors by using the commercially available AMP-Glo^TM^ kit. Our HTS assay demonstrated high robustness, meeting the NIH HTS Assay Validation guidelines’ acceptance criteria: a Z’-factor ≥ 0.4 or a signal window (SW) ≥ 2, and a coefficient of variation (CV) ≤ 20%^40^. We also present the results of a pilot high-throughput screening of a Core compound library from the Arizona Center for Drug Discovery (ACDD), comprising 5,120 blood-brain-barrier (BBB)-permeable compounds across sixteen 384-well plates. Several novel inhibitors of UBA5 were identified and characterized by using biochemical, biophysical, and cellular assays.

## Results

### AMP-Glo^TM^-based HTS assay

To develop an *in vitro* UFMylation assay suitable for HTS, we performed codon optimization, synthesized, cloned, expressed and purified UFM1 and UBA5_37-346_ encoding UBA5 amino acids (aa) 37-346, representing the functional fragment of UBA5^41–46^.

We initially explored a few different assay platforms. In the UFM1 activating reaction catalyzed by UBA5 (UFM1+UBA5+ATP →UFM1-UBA5+AMP+PPi) (**Figure 1A**), ATP is converted to AMP and pyrophosphate (PPi), and UFM1 is covalently conjugated to UBA5 (UFM1-UBA5) through several intermediate steps^37^. Our initial efforts to use commercial kits to quantify ATP depletion or to quantify PPi production were unsuccessful. For reasons unknown, the reagents from the PPi kits from commercial vendors produced high background readings when mixed with UBA5 protein alone, making it impossible to develop a meaningful HTS assay. Additionally, attempts to directly quantify ATP depletion using the Promega Kinase-Glo^TM^ kit were also unsuccessful due to a low signal-to-background (S/B) ratio of only approximately 1.5-fold.

**Figure 1.**
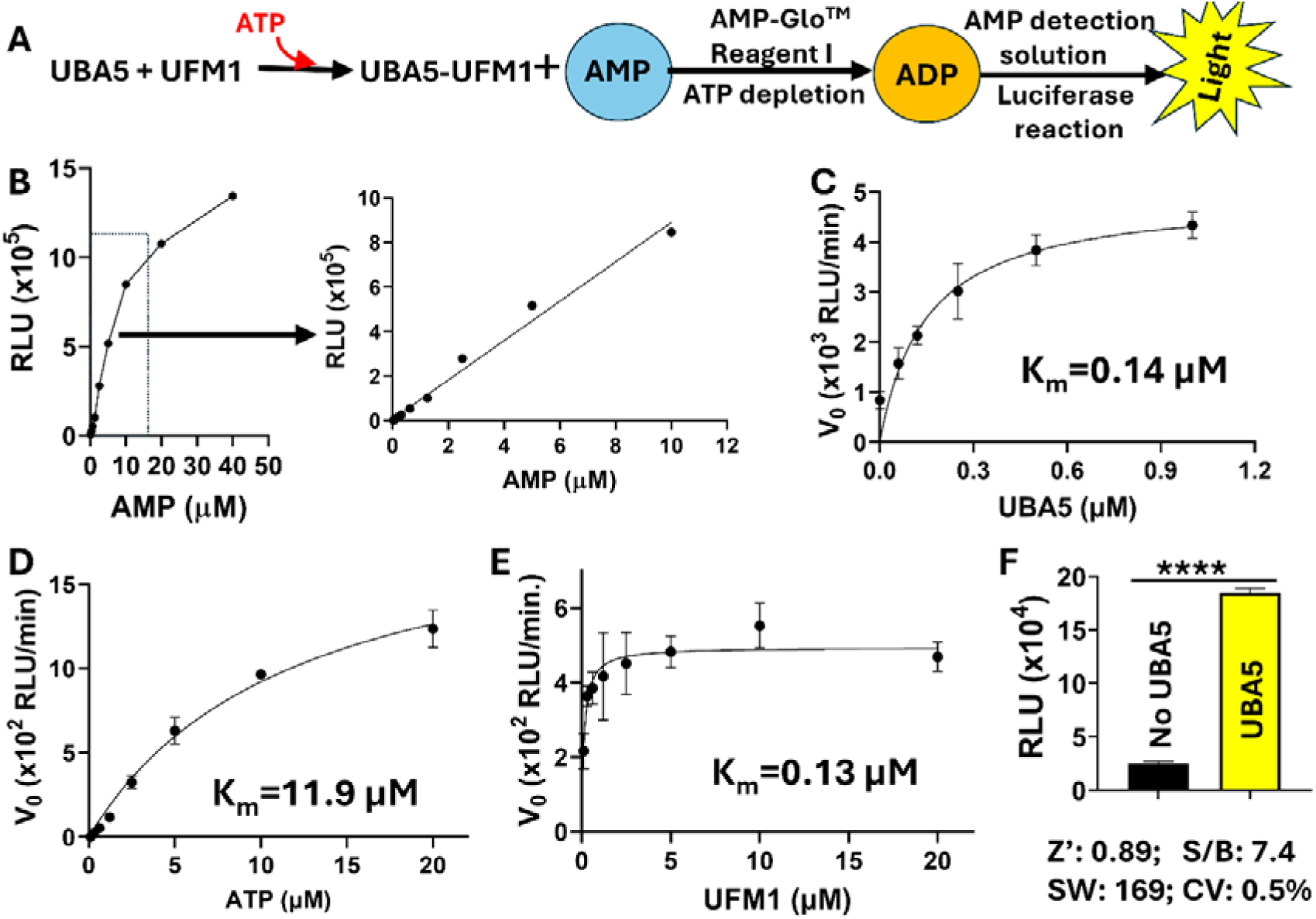
Development and optimization of AMP-Glo^TM^-based UBA5 UFMylation assay. (**A**) Assay principle. (**B-E**) Kinetic analyses of the AMP-Glo^TM^-based UBA5 coupling assay to optimize assay parameters and assay components including AMP (**B**), UBA5 enzyme (**C**), ATP (**D**) and UFM1 substrate (**E**), using 384-well plate. Each experiment varied only the indicated component in a concentration series, while all other components were kept at fixed concentrations. RLU, relative fluorescence unit. (**F**) HTS parameters for the optimized AMP-Glo^TM^ UBA5 assay in a 384-well plate, by comparing the RFU generated with and without UBA5. N=16. ****, p<0.0001. For all AMP-Glo assays here and below (unless otherwise specified): UBA5, 0.8 µM; UFM1, 5 µM; ATP, 5 µM.

Finally, we developed a luminescence-based coupling UBA5 assay to quantify AMP production by using the Promega AMP-Glo^TM^ kit (**Figure 1A**). The kit’s Reagent I first depletes the input ATP and converts AMP to ADP. Then, Reagent II converts ADP back to ATP, which is detected by the firefly luciferase in the kit, generating luminescence. Thus, the UFM1-activation reaction catalyzed by UBA5 can be monitored by measuring the luminescence.

We first established the linear detection range of AMP using the kit, ensuring that the generated luminescence corresponds directly and linearly to AMP production (**Figure 1A, 1B**). This step is crucial to guarantee that the inhibitor’s effectiveness will have a linear relationship with the luminescence generated. As shown in **Figures 1A** and **1B**, AMP concentrations within the 0–10 µM range showed a linear relationship with luminescence production. The addition of up to 100 µM ATP did not significantly alter this trend or the signal (data not shown), indicating that Reagent I in the kit effectively depleted the input ATP up to 100 µM without any issues. Next, we conducted kinetic analyses to optimize the assay, determining the ideal concentrations of ATP, UBA5_37-346_, and UFM1, as well as optimizing buffer conditions, DMSO and detergent tolerance, and incubation time (**Figure 1C-1E**). These adjustments were made to ensure the reaction remained within a linear range and achieved an optimal signal-to-background (S/B) ratio as well as a more appropriate HTS parameter: signal window (SW), in accordance with the NIH HTS Assay Validation guide^40^. As shown in **Figure 1C**, dose-dependent increase of luminescence was observed by increasing UBA5 concentration. The result indicates that UBA5 is active, catalyzes UFM1 conjugation to UBA5, and converts ATP to AMP, leading to luminescence increase. By fitting the kinetic data, a *K_m_* of 0.14 µM was obtained (**Figure 1C**).

Similarly, kinetic studies were also performed for UFM1 and ATP in concentration series (**Figure 1D, 1E**). As concentrations of UFM1 and ATP increased, hyperbolic increases in luminescence were obtained. *K_m_* values of 11.9 µM for ATP and 0.13 µM for UFM1 were determined by fitting the kinetic data.

Upon optimization, we achieved superior HTS parameters by comparing endpoint luminescence readings of reactions with and without UBA5, resulting in a Z’ factor of 0.89, S/B ratio of 7.4, SW of 169, and CV of 0.5% (**Figure 1F**).

### Inhibition of UBA5 activity by positive control inhibitor DKM 2-93

We next evaluated the UBA5 inhibitor DKM 2-93 (DKM), previously reported in the literature^39^, using the AMP-Glo^TM^ coupling assay. Using the AMP-Glo^TM^ assay, we showed that DKM dose-dependently inhibited UBA5 activity, with an *IC_50-UBA5-Glo_* value of 547 µM (**Figure 2A**), which is comparable to the previously reported *IC_50_* of 430 µM^39^. Using DKM as a control compound, we demonstrated the robustness of the AMP-Glo^TM^ UBA5 coupling assay in a 384-well plate format, achieving a satisfactory Z’ factor of 0.85, S/B ratio of 5.8, SW of 37, and CV of 2% against UBA5 (**Figure 2B**). The assay showed excellent well-to-well consistency (**Figure 2C**). These parameters were reproducible over multiple plates and dates (**Figure 2D**). We did not observe drifts or edge effects with this assay (data not shown).

**Figure 2.**
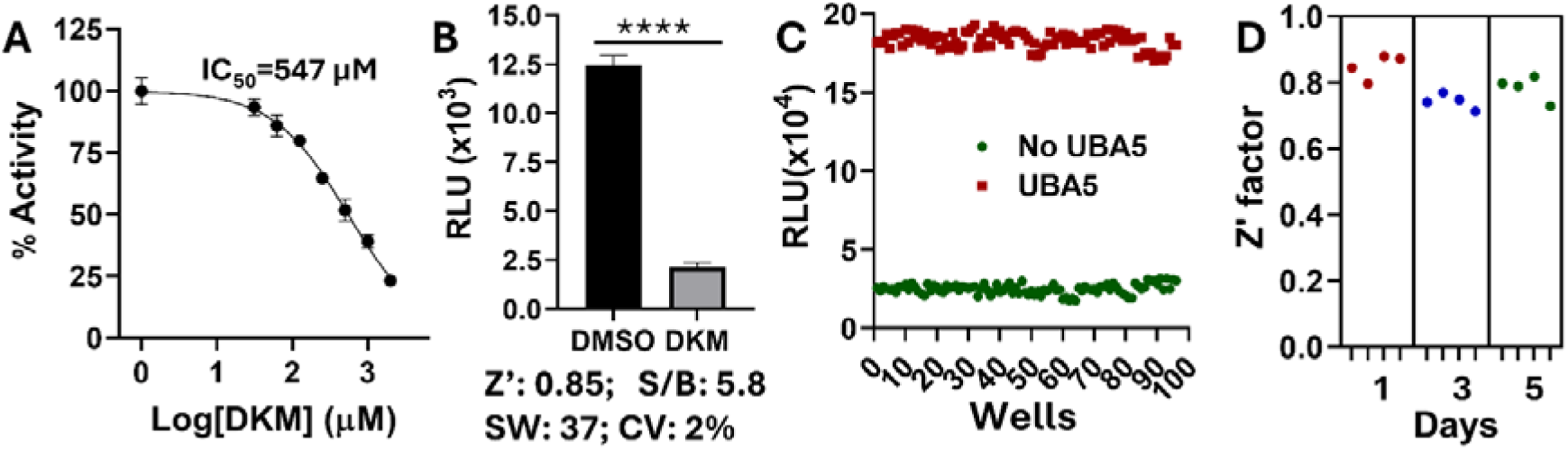
HTS AMP-Glo^TM^ UBA5 assay. (**A**) Dose response curve of inhibition of the UBA5 activity by concentration series of DKM 2-93. N=3. Data were normalized using DMSO-treated wells with UBA5 set as 100%, and wells without UBA5 set as 0%. (**B**) DKM (3 mM) inhibited UBA5 activity in a 384-well plate. N=96. ****, P<0.0001. (**C**) Well-to-well relative luminescence in 384-well format with and without UBA5; N=96. (**D**) Z’ factor over plates and dates. Each spot represents Z’ score calculated from the results of 384-well plates on 4 different plates/day on 3 dates.

### Pilot HTS

Next, we conducted a pilot HTS against a Core compound library of the Arizona Center for Drug Discovery, consisting of 5,120 blood-brain-barrier (BBB)-permeable compounds across sixteen 384-well plates. DKM served as a positive control inhibitor. The results demonstrated the robustness of the AMP-Glo^TM^ UBA5 assay, with an average Z’ score of 0.79, S/B ratio of 6.4, SW of 58, CV of 5.3% (**Figure 3A**). These metrics met the acceptance criteria for a reliable HTS assay, which are Z’ ≥ 0.4, SW ≥ 2, and CV ≤ 20%, according to the NIH HTS Assay Validation guidelines^40^.

**Figure 3.**
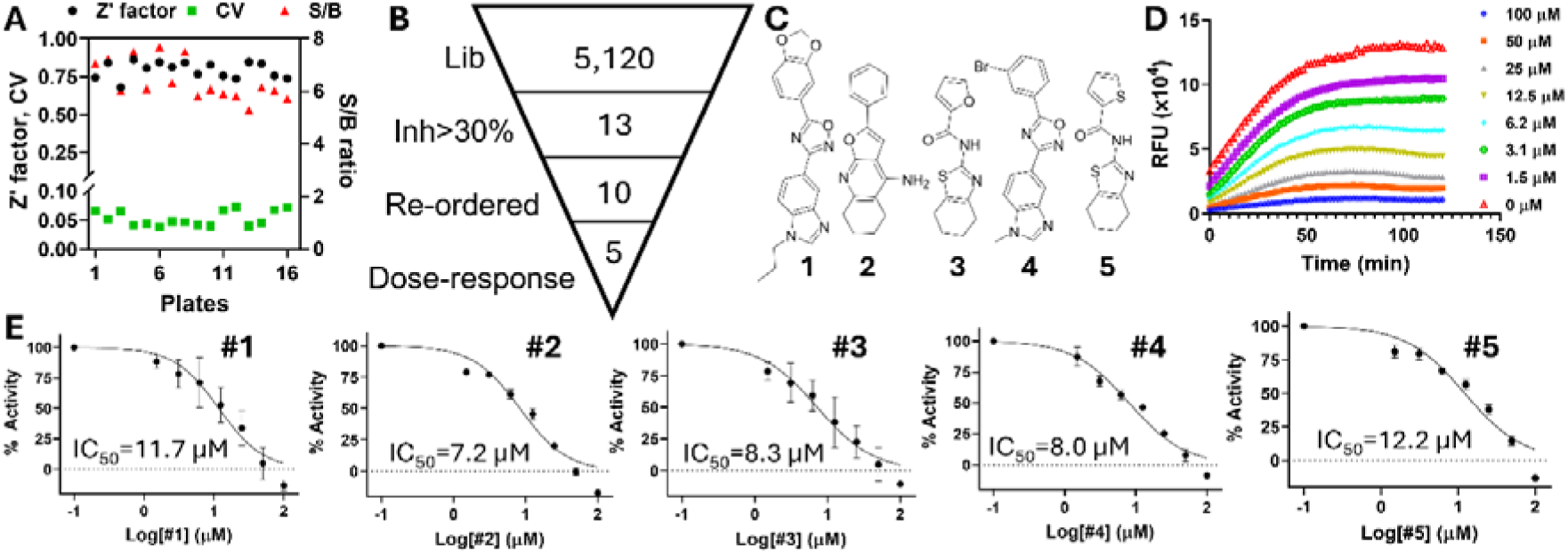
Pilot HTS. (**A**) Z’ factor, S/B ratio, and CV over plates of a pilot HTS against a CNS compound library in sixteen 384-well plates. (**B**) Screening flowchart. (**C**) Compound structures of hits. (**D**) Representative kinetic readings of dose-response inhibition of UBA5 by Comp#**2**. (**E**) Dose response inhibition of UBA5 activity by 5 hit compounds. Data were normalized using DMSO-treated wells with UBA5 set as 100%, and wells without UBA5 set as 0%. N=3. We identified 13 compounds at 20 µM that showed significant inhibition (>30%), yielding a hit rate of 0.25% (Figure 3B**, 3C**). After cheminformatic analysis, three hits were removed as pan-assay interference compounds (PAINS)^47,48^, leaving 10 compounds. Powders for these remaining compounds were re-ordered for further analysis.

Confirmation experiments revealed that 5 out 10 compounds showed dose-dependent inhibition of UBA5 activity with *IC_50-UBA5-Glo_* values in low micromolar range (**Figure 3C-3E, Table 1**). Notably, all identified hits possess low molecular weight (<400 Da), which is a good size for the hit-to-lead discovery, and have CNS multiparameter optimization (MPO) scores greater than 4.0, which is the desired threshold for compounds suitable for treating CNS diseases^49^. The five hits identified possess three distinct scaffolds. Hits **1** and **4** contain “1,2,4-oxadiazole” ring as the central piece flanked by a fused bicycle – “1-alkyl-1*H*-benzo[*d*]imidazol-5-yl-“ – and a phenyl ring in 3 and 5 position, hit **2** possesses a fused tricyclic “tetrahydronaphtho[2,3-*b*]furan-4-amine” core substituted with a phenyl ring, and **3** and **5** are amide derivatives prepared with a common fused bicyclic amine, 4,5,6,7-tetrahydrobenzo[*b*]thiophen-2-yl-amine, with 2-furan- and thiofuran carboxylic acids. The structural features of all 5 hits suggest that these may or may not occupy the same site of the enzyme or the coupling enzymes, especially as **1**, **3**, **4**, and **5** are likely to form a curved shape, whereas hit **2** seems to form a rod-like structure. All hits could potentially generate sound hydrophobic contacts and hydrogen bonding(s) through various rings and the hydrogen donating and accepting groups.

**Table 1.** Compounds properties.

| | *MW*<br>(Da) | *LogD* | *MPO* | *TPSA*<br>(Å) | *IC*<sub>50-UBA5-Glo</sub><br>($\mu\text{M}$) | *IC*<sub>50-UBA5-FP</sub><br>($\mu\text{M}$) | *IC*<sub>50-UBA1-Glo</sub><br>($\mu\text{M}$) | *SI*<sub>El</sub> | *IC*<sub>50-coupling-Glo</sub><br>($\mu\text{M}$) | *SI*<sub>Coupling</sub> | *CC*<sub>50</sub><br>($\mu\text{M}$) |
| --- | --- | --- | --- | --- | --- | --- | --- | --- | --- | --- | --- |
| **1** | 348.36 | 3 | 4.8 | 75.2 | 11.7 | 16.6 | >100 (42%*) | >8.5 | 23.7 | 2.0 | >200 |
| **2** | 264.33 | 2.1 | 4.8 | 52 | 7.2 | 11.7 | 80 (54%*) | 11 | 8.6 | 1.2 | 168 |
| **3** | 248.31 | 3.2 | 5 | 83.4 | 8.3 | 14.0 | 95 (57%*) | 11 | 51.5 | 6.2 | >200 |
| **4** | 355.19 | 4 | 4.2 | 56.7 | 8.0 | 9.7 | >100 (18%*) | >12 | 35.1 | 4.4 | >200 |
| **5** | 264.38 | 3.3 | 4.8 | 98.5 | 12.2 | 13.1 | 59 (73%*) | 4.8 | 95.6 | 7.8 | >200 |

Note: MW, molecular weight; SI, selectivity index; $SI_{El} = IC_{50-UBA1-Glo} / IC_{50-UBA5-Glo}$ ; $SI_{Coupling} = IC_{50-coupling-Glo} / IC_{50-UBA5-Glo}$ ; \*, percent of inhibition at 100 $\mu\text{M}$ .

### Development of a secondary assay based on fluorescence polarization (FP)

False positives may arise from compound aggregation, PAINS properties, or interference with enzymes in the coupling assay. Therefore, developing secondary or tertiary assays to prioritize hit compounds is essential. To address this, we developed a secondary assay using fluorescence polarization (FP) with the Transcreener AMP²/GMP² FP kit (Bellbrook Labs) (**Figure 4A**). This assay helps further characterize UBA5 activity by monitoring AMP production through FP changes and allows us to further prioritize the hit compounds.

**Figure 4.**
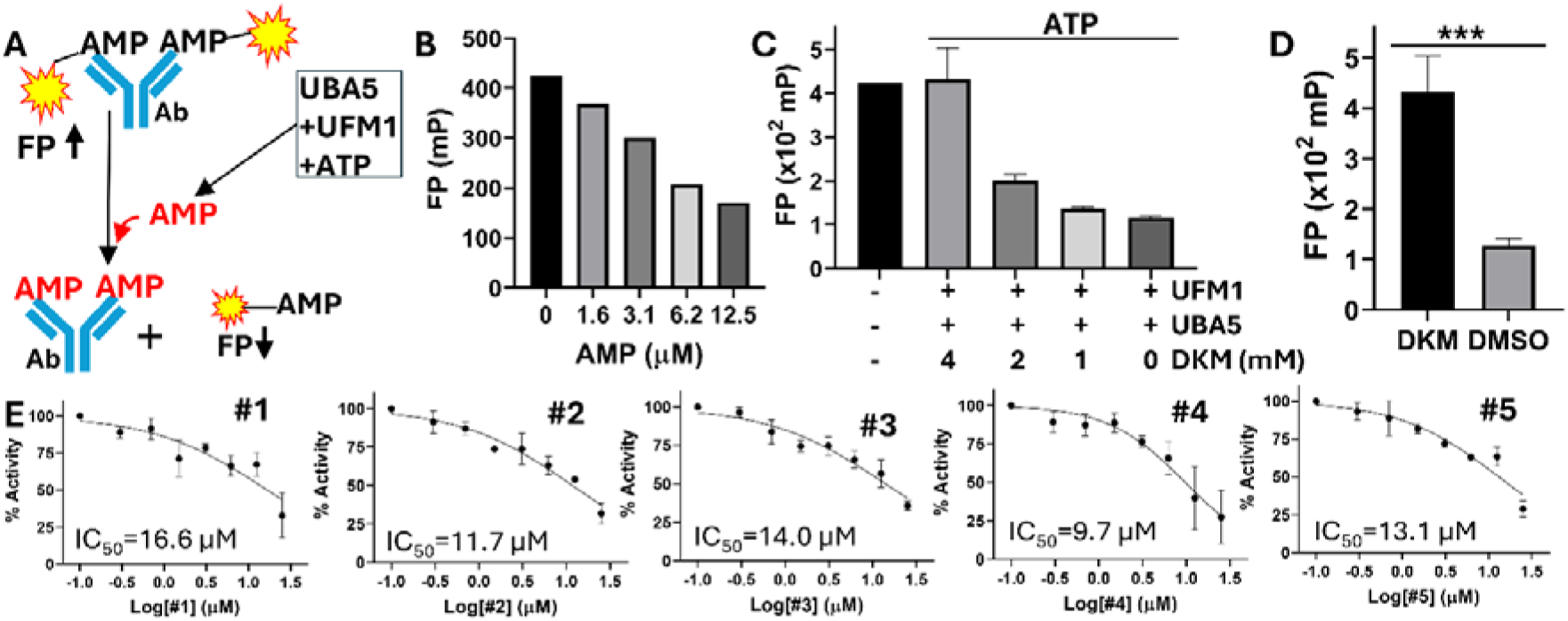
Secondary FP-based assay. (**A**) Principle of FP-based secondary assay. (**B**) AMP titration using the FP assay based on the Transcreener® AMP/GMP FP Assay Kit (Bellbrook), following the manufactory protocol. (**C**) DKM titration against UBA5 using the FP assay. N=3. For all FP assays here and below (unless specified): UBA5, 1.0 µM; UFM1, 2.5 µM; ATP, 20 µM. (**D**) DKM (4 mM) inhibited UBA5 activity using the FP assay in a 384-well plate. N=16. ***, P<0.001. (**E**) Fitting of dose-response curve of inhibition of UBA5 by compounds, using the FP-based secondary assay. Data were normalized using DMSO-treated wells with UBA5 set as 100%, and wells without UBA5 set as 0%. N=3.

In the FP assay, an AlexFluor-633-labeled AMP^2^/GMP^2^ tracer (AMP-633) binds an antibody (Ab) specially recognizing AMP or GMP, resulting in high FP values (**Figure 4A**). However, when free AMP is produced by the UBA5+UFM1 reaction, it displaces the AMP-633 tracer from the antibody, leading to a decrease in FP. In the presence of a UBA5 inhibitor, AMP production is diminished, resulting in less displacement of the AMP-633 tracer and maintaining high FP values.

As shown in **Figure 4B**, we initially observed that AMP dose-dependently reduced FP, demonstrating that free AMP effectively displaced the AMP-633 tracer from the antibody in a dose-dependent manner. Next, we compared UFMylation reactions with and without UBA5. As shown in **Figure 4C**, the presence of UBA5 in the reaction (last column) significantly reduced FP values compared to the reaction without UBA5 (1^st^ column). These results suggest that UBA5 reacts with UFM1, converting ATP to AMP, which then displaces the AMP-633 tracer from the antibody, leading to a low FP value.

Further, we used DKM as a control inhibitor to optimize the FP-based assay parameters for monitoring the UBA5-UFM1 reaction, following a similar approach to the optimization of the AMP-Glo^TM^ assay (data not shown). Under optimized conditions, DKM dose-dependently inhibited the UBA5-mediated UFMylation reaction (**Figure 4C**). The assay proved to be robust, with a Z’-factor of 0.5, an S/B ratio of 3.7, an SW of 5.7, and a CV of 8.2%, meeting the NIH HTS Assay Validation guidelines^40^ (**Figure 4D**).

We then used this FP-based assay as a secondary assay to characterize compounds identified from the primary AMP-Glo^TM^ screening. The results showed that all five compounds dose-dependently inhibited UBA5 activity, with *IC_50-UBA5-FP_* values falling within a similar micromolar range (**Figure 4E**, **Table 1**). The slight discrepancy between *IC_50_* values measured using AMP-Glo^TM^ and FP assays may be due to assay sensitivity, with the FP assay being slightly less sensitive, requiring a slightly higher UBA5 concentration than the AMP-Glo^TM^ assay.

### UBA5-free coupling enzyme assay

To determine if the identified hits inhibit the coupling enzymes in the AMP-Glo^TM^ kit, we performed a UBA5-free coupling enzyme assay. In this assay, AMP served as the direct substrate for the coupling enzymes in the AMP-Glo^TM^ kit, along with all other components such as ATP, UFM1, and the reagents from the coupling kit, but without UBA5. The UBA5-free AMP-Glo^TM^ assay demonstrated sensitivity comparable to the primary AMP-Glo^TM^ assay in the presence of UBA5 (**Figure 5A**). When AMP was added to the kit, it directly generated significant luminescence compared to the reaction without AMP, resulting in a Z’ factor of 0.81, an S/B ratio of 17.7, an SW of 37.0, and a CV of 4.1%. These parameters met the NIH HTS Assay Validation guidelines^40^.

**Figure 5.**
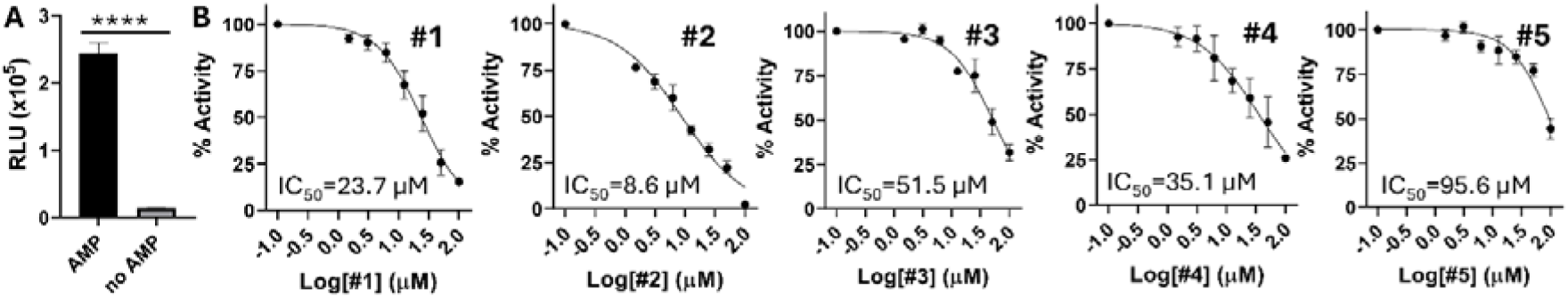
UBA5-free AMP-Glo^TM^ assay. (**A**) AMP-Glo assay in the absence of UBA5. AMP (5 µM) was used as a direct substrate. UFM1, 5 µM. (**B**) Fitting of dose-response curve of inhibition of the coupling enzymes in the AMP-Glo kit by compounds, using the coupling enzyme tertiary assay. AMP, 5 µM; UFM1, 5 µM. Data were normalized using DMSO-treated wells with AMP set as 100%, and wells without AMP set as 0%. N=3.

We next used this UBA5-free AMP-Glo^TM^ coupling enzyme assay to assess the identified compounds. As shown (**Figure 5B**, **Table 1**), except for Comp **2**, which exhibited strong inhibition of the coupling enzymes in the kit, the other four hits only moderately inhibited the enzymes in the kit. Their *IC_50-coup-Glo_* values were significantly—by at least 2.0 to 7.8-fold—higher than their corresponding *IC_50-UBA5-Glo_* values, indicating that their primary inhibitory activity targets UBA5 rather than the coupling enzymes in the kit.

### Selectivity assay

UFM1 is a member of the ubiquitin-like protein (UBL) family, which is covalently attached to target proteins to regulate their activities. This process occurs *via* a general E1-E2-E3 multienzyme cascade. A total of 17 human UBLs from 9 phylogenetic classes have been identified as being conjugated to various molecules^50,51^. While all UBLs share a similar overall structural fold, each UBL typically operates through its own unique E1–E2–E3 enzyme cascade and exerts specific effects on its respective targets. In humans, eight E1 enzymes—UBA1, UBA2/SAE1, UBA3/NAE1, UBA4-7, and ATG7—have been identified as key initiators responsible for the conjugation of specific UBLs^50,51^. E1 enzymes are divided into canonical and noncanonical families. Canonical E1 enzymes, such as UBA1, UBA2/SAE1, UBA3/NAE1, UBA6, and UBA7, are responsible for activating Ub, the SUMO protein family, NEDD8, FAT10, and ISG15, respectively. They possess two pseudosymmetric adenylation domains, which are encoded by either one or two genes. In contrast, noncanonical E1 enzymes, such as ATG7, UBA4, and UBA5, which respectively activate the ATG8 and ATG12 protein families, URM1, and UFM1, form homodimers to perform the E1 function. Despite having distinct structures, all E1s share a common catalytic mechanism, which involves the conversion of ATP to AMP during the UBL activation process^50,51^. Therefore, identifying inhibitors that are specific to UBA5, without affecting other E1 enzymes, is crucial.

Given the shared ATP-to-AMP conversion mechanism among E1s, we hypothesized that the AMP-Glo^TM^ assay could be adapted for use with other E1s as a specificity assay for identified hits. To demonstrate this, we cloned, expressed, and purified His-tagged UBA1 in a manner similar to UBA5. The AMP-Glo assay successfully detected UBA1 activity (**Figure 6A**). In the presence of UBA1, we observed a 6.6-fold increase in luminescence compared to the control without E1, indicating that UBA1, like UBA5 in its UFMylation reaction, converts ATP to AMP during Ub activation, leading to luminescence increase (**Figure 6A**). We optimized the assay conditions similarly to those for UBA5, observing a dose-dependent increase in luminescence against ATP, UBA1 and other components (**Figure 6B**). Using this specificity assay, we found that these hits only moderately inhibited UBA1 (**Table 1**, **Figure 6C,6D**). Specifically, compounds **1** and **4** exhibited *IC_50-UBA1-Glo_* values larger than 100 µM, while compounds **2**, **3** and **5** showed moderate *IC_50-UBA1-Glo_* values between 59 to 95 µM. These values are 4.8-to 12-fold higher than those for UBA5 (**Table 1, *SI_E1_***), suggesting that the hits selectively target UBA5.

**Figure 6.**
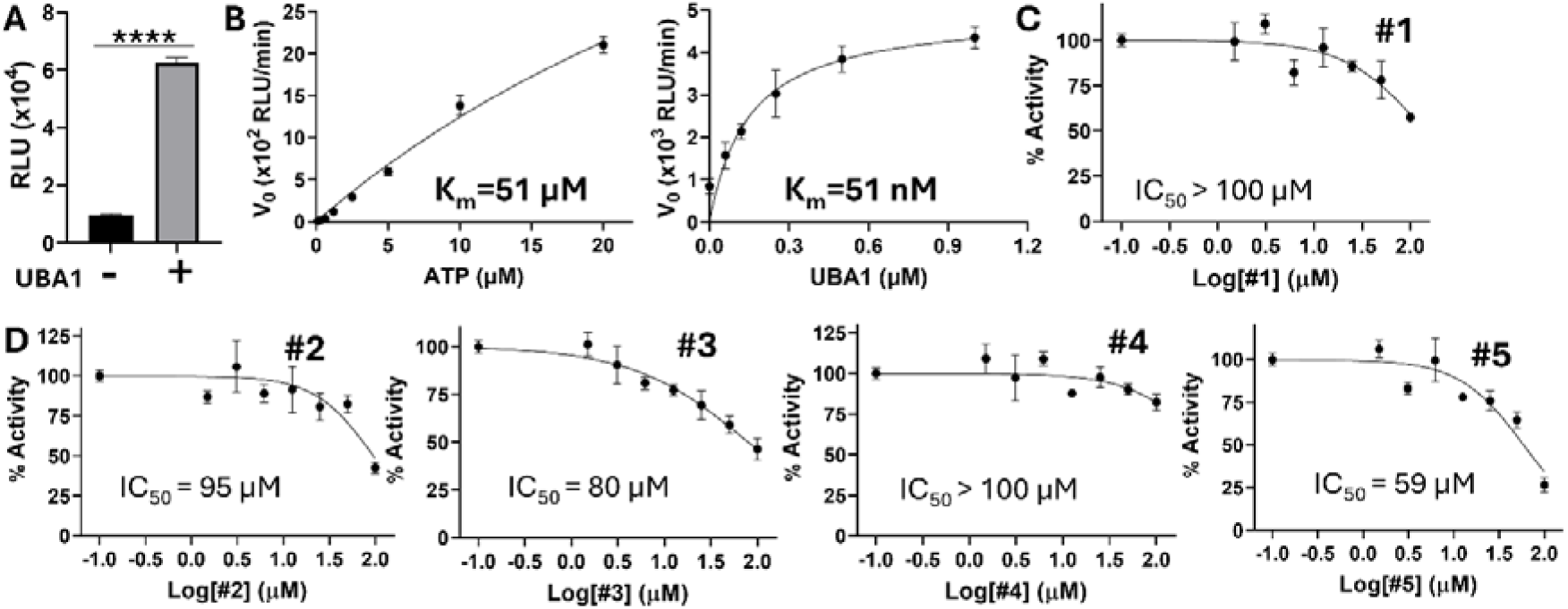
Quaternary E1 specificity assay. (**A**) UBA1 AMP-Glo^TM^ specificity assay with and without UBA1. For all UBA1 AMP-Glo assays here and below (unless specified): UBA1, 0.3 µM; Ub, 10 µM; ATP, 5 µM. ****, p<0.0001. (**B**) Kinetic analysis of ATP and UBA1 titration for the UBA1 AMP-Glo specificity assay using 384-well plate. (**C-D**) Titration of identified Comps **#1-5** against UBA1. Data were normalized using DMSO-treated wells with UBA1 set as 100%, and wells without UBA1 set as 0%. N=3.

### Identified hits inhibited UFMylation *in vitro* in gel-based assay

We next evaluated if the hits inhibited UBA5-UFM1 conjugation *in vitro* using a gel-based assay. Our results are consistent with those found using AMP-Glo^TM^ or FP-based assays. All hits dose-dependently inhibited UFM1 charging to UBA5 (**Figure 7**).

**Figure 7.**
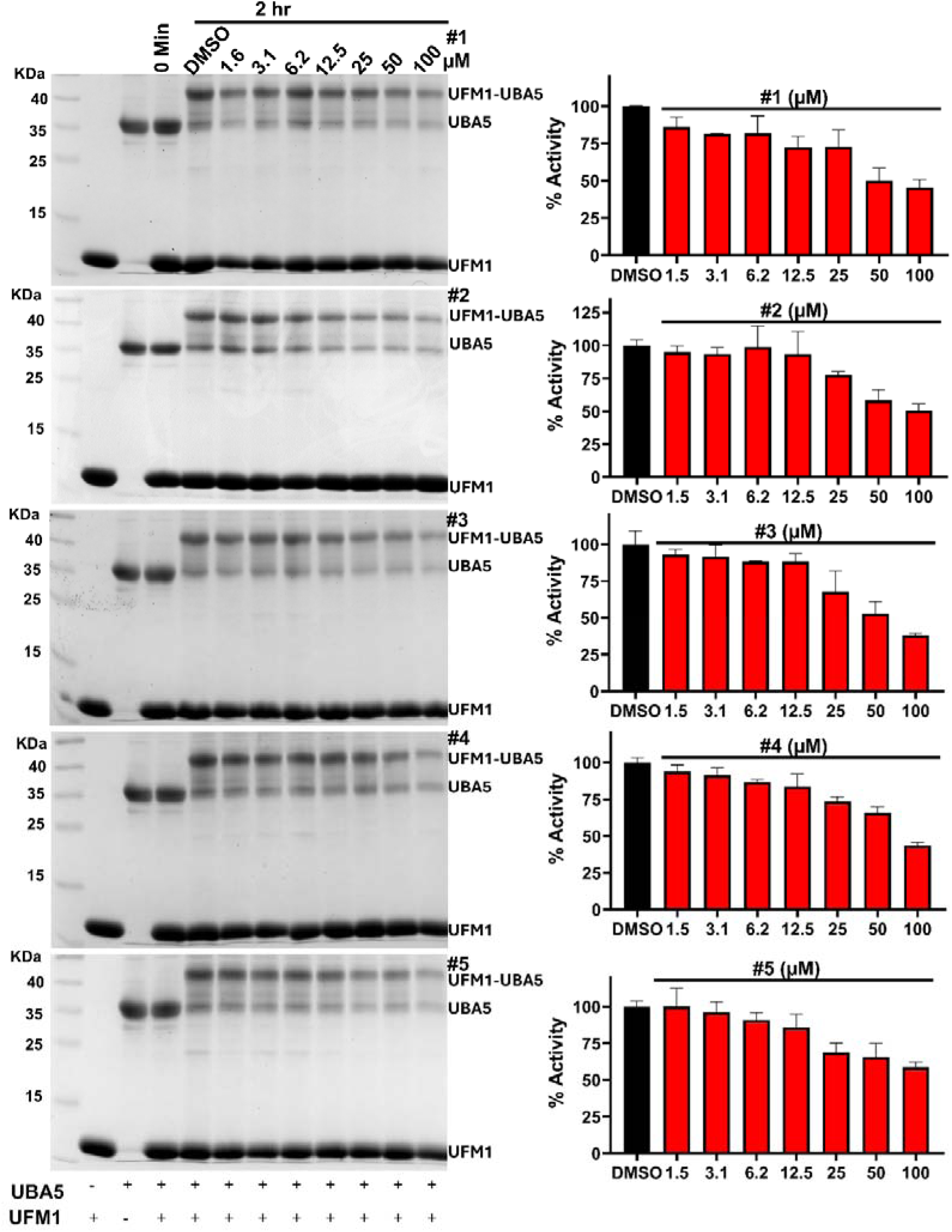
Inhibition of UFMylation by hit compounds *in vitro*. Inhibition of UFMylation of UBA5 was assessed using an *in vitro* in-gel-based UFMylation assay (left panel), with accompanying band quantification (right panel). UBA5, 12 µM; UFM1, 25 µM; ATP, 2 mM. Samples were resolved on 12% SDS-PAGE gels and stained with Coomassie Blue. Band intensities were quantified using ImageJ. Data were normalized by calculating the ratio of the band intensity of the UFM1-UBA5 complex to the combined intensity of UFM1 and UBA5 alone, with the average of the DMSO-treated wells at the 2-hour time point set as 100% and those at the 0-hour time point set as 0%. N=2.

### Inhibited UFMylation in cell-based UFMylation assay

Subsequently, we selected three representative compounds—compounds **1**, **2**, and **3**, each from a different scaffold class—to evaluate their ability to inhibit the endogenous UFMylation process in HEK293T cells. After treating the cells with the compounds, we assessed the UFMylation levels of UBA5 and its downstream E2, UFC1, using an anti-UFM1 antibody (**Figure 8**). Results indicated that all three compounds inhibited endogenous UFMylation of both UBA5 and UFC1 in a dose-dependent manner. Notably, at 60 µM, compound **3** reduced UFMylation of both UBA5 and UFC1 by nearly 50%.

**Figure 8.**
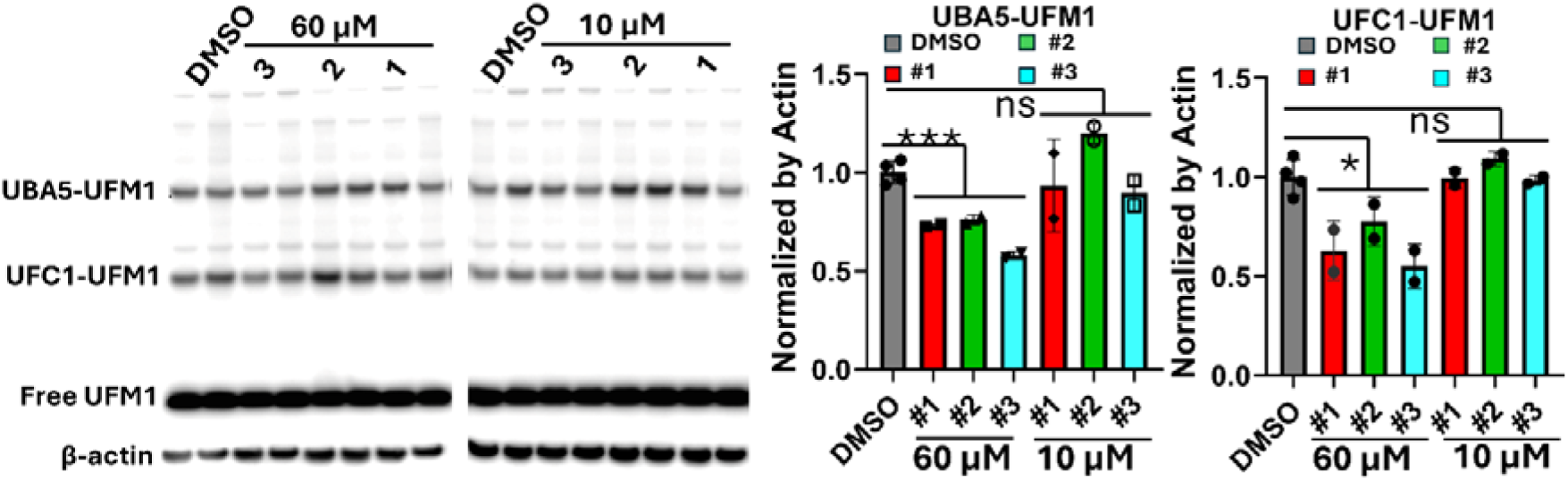
Inhibition of UFMylation of UBA5 and UFC1 by representative compounds in cell. Inhibition of UFMylation of UBA5 was assessed using a cell-based UFMylation assay (left panel), with accompanying band quantification (right panel). Briefly, HEK293T cells were seeded in 12-well plates, cultured to approximately 80% confluent, and incubated with compounds at concentrations of 10 µM and 60 µM for 24 hours. After incubation, the cell media were removed, and cells were lysed with 100 µL RIPA lysis buffer. Protein concentrations were quantified using the BCA assay, and western blotting was conducted with an anti-UFM1 antibody on a 4-12% Bis-Tris Gel (Invitrogen) (left panel). The UBA5-UFM1 and UFC1-UFM1 bands were quantified using ImageJ (right panel), with band intensities normalized to β-actin. The DMSO-treated sample average was set to 1.0 as the baseline for comparison across treatment groups. N=2 (treatment groups). N=4 (DMSO). ***, p<0.001; *, p<0.05.

### Cytotoxicity analysis

We next evaluated the cell cytotoxicity of these compounds using a WST-8 cell viability assay as we described previously^52–55^(**Table 1**, **Figure 9**). The results indicate that none of the hits display cytotoxicity (*CC_50_*) to astrocytes-like CCF-STTG1 cells. (*CC_50_*: compound concentration required to reduce cell viability by 50%).

**Figure 9.**
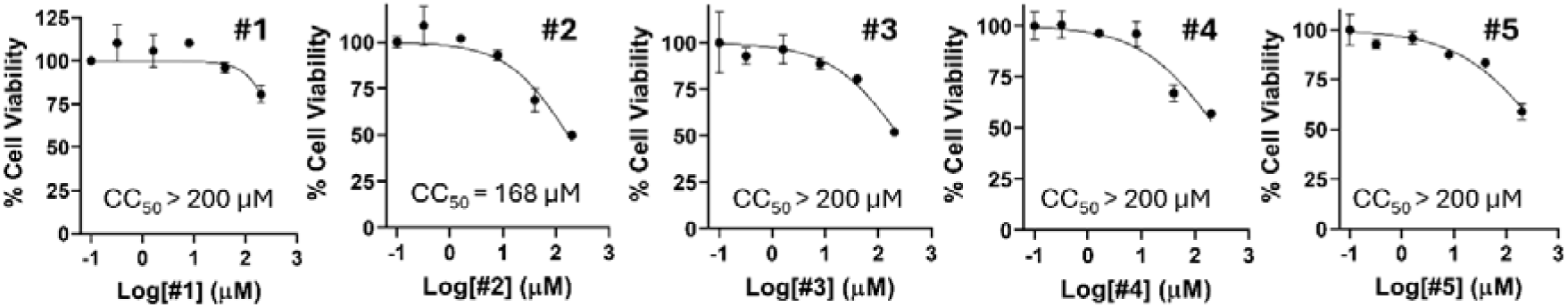
Cell viability against astrocyte-like CCF-STTG1 cells. CCF-STTG1 (ATCC) cells were incubated with various concentrations of compounds, and then viability was assayed at 48 h of incubation, using the WST assay. Data were normalized by setting the absorption readings of cells treated with DMSO to 100% and the readings from cell-free wells treated with DMSO to 0%. N = 3.

## Discussion

UBA5 plays a crucial role in the UFMylation pathway, a post-translational modification process. There is growing evidence linking the dysregulation of UFMylation to various human diseases, including neurodegenerative disorders and cancers^4,7–19^. Given its central role in cellular homeostasis, targeting UBA5 with specific inhibitors presents a promising therapeutic strategy.

In this study, we developed an HTS assay to identify novel inhibitors of UBA5, the E1-activating enzyme critical to the UFMylation cascade. Our HTS assay, based on the AMP-Glo™ kit (Promega), demonstrated robustness and reproducibility, allowing the successful screening of a small BBB-permeable compound library. Through this screening, we identified several novel UBA5 inhibitors, which were further characterized by secondary biochemical, biophysical, and cellular assays.

One key finding was the identification of five novel UBA5 inhibitors, belonging to three distinct chemical scaffolds, with low micromolar *IC_50_* values. Importantly, these compounds demonstrated good selectivity for UBA5 over other E1 enzymes, including UBA1, which activates ubiquitin, underscoring their potential for selective targeting of the UFMylation pathway. Selectivity is crucial since many E1 enzymes share common structural and mechanistic features^50,51^, making off-target effects a significant concern in the development of E1 inhibitors.

Moreover, these inhibitors showed promising activity in both *in vitro* assays and cellular models. Four of the five compounds effectively inhibited endogenous UFMylation in HEK293T cells, reducing the conjugation of UFM1 to UBA5 and UFC1, the downstream E2 enzyme in the pathway. These findings suggest that our identified inhibitors can effectively suppress UFMylation in a cellular context, which is critical for their potential therapeutic development.

The ability to target UBA5 in diseases like Alzheimer’s and cancer could open new avenues for therapy. UBA5 has been implicated in protein homeostasis and stress response, both of which are crucial in neurodegenerative diseases^1–3,20–23,25–30^. Additionally, aberrant UFMylation has been linked to oncogenesis, with UBA5 overexpression associated with tumor progression and poor prognosis in multiple cancer types^4,11,31–36^. Our work supports the hypothesis that inhibiting UBA5 could modulate these pathways, providing therapeutic benefits.

Despite the promising results, there are still challenges and limitations to address. While our hits demonstrated reasonable selectivity for UBA5, further optimization is necessary to improve their potency and pharmacokinetic properties. Structural studies, including co-crystallization with UBA5, could provide insight into the binding modes of these inhibitors and guide rational drug design for enhanced efficacy. Additionally, detailed *in vivo* studies are needed to evaluate the therapeutic potential of these inhibitors in disease models, particularly neurodegenerative and cancerous conditions where UFMylation dysregulation plays a key role.

### Conclusion

This study demonstrates the successful development of an HTS assay for identifying UBA5 inhibitors and the discovery of several promising candidates with selective inhibitory activity. These findings provide valuable tools for studying UFMylation and lay the groundwork for developing potential therapies targeting UBA5 in diseases such as Alzheimer’s and cancer. Ongoing research into the identification and characterization of UBA5 inhibitors will offer crucial insights into their potential clinical applications. Further structural and pharmacological optimization of these inhibitors could lead to the development of clinically relevant compounds, opening new therapeutic avenues for modulating UFMylation in human disease.

## MATERIALS AND METHODS

### Materials

Ubiquitin was purchased from South Bay Bio (cat# SBB-UP0013). Anti-UFM1 antibody was purchased from Abcam (cat# 109305).

### Expression and purification of UFM1, UBA1 and UBA5_37-346_

The codon-optimized gene sequence of UFM1, including a TEV protease-recognition site at the N-terminus, was synthesized and inserted between the His-tag and EcoR1 site in the pET28a vector using seamless cloning technology by GeneUniversal. The expression plasmid UFM1-p28 was transformed into *E. coli* Rosetta (DE3) cells, which were then cultured in Luria broth medium supplemented with 100 μg/mL kanamycin at 37 °C until the optical density (OD) at 600 nm reached 0.6. The cells were induced with 0.3 mM IPTG and further incubated with shaking at 150 rpm and 16 °C for 18 hours. After incubation, the cells were collected by centrifugation at 8000 rpm for 20 minutes. The cell pellets were resuspended in lysis buffer (20 mM HEPES, pH 7.5, 200 mM NaCl), and lysozyme along with phenyl methyl sulfonyl fluoride (PMSF) was added. Cells were lysed using sonication and then centrifuged at 15,000 rpm at 4 °C for 30 minutes. The supernatant was loaded onto a pre-charged Ni-NTA affinity column (Qiagen) and washed with the resuspension buffer containing 30 mM imidazole. His-tagged UFM1 was eluted using lysis buffer containing 50 mM and 100 mM imidazole. The UFM1 was further purified by size-exclusion chromatography using a 75 Superdex column, and peak fractions were collected and pooled. The purified UFM1 was stored in a buffer containing 25 mM HEPES, pH 7.5, 150 mM NaCl, and 2 mM DTT.

The codon-optimized gene sequence of UBA5 encoding UBA5 amino acids 37-346 was synthesized and cloned into a custom His-SUMO vector using seamless cloning technology by GeneUniversal. The expression and purification of UBA53_7−346_-His-SUMO were performed similarly to UFM1, with the exception of the on-column digestion step. The supernatant was loaded onto a pre-charged Ni-NTA affinity column (Qiagen). On-column digestion was carried out using the ULP1 protease, with incubation for 2 hours at room temperature (RT). Following digestion, flowthrough fractions were collected, washed with 2 columns of the resuspension buffer, and combined. The UBA5_37−346_ was then concentrated and further purified by size-exclusion chromatography using a 75 Superdex column. Peak fractions were collected and pooled, and the purified UBA5 was stored in a buffer containing 25 mM HEPES, pH 7.5, 150 mM NaCl, and 2 mM DTT.

The codon-optimized gene sequence of UBA1 was synthesized and cloned into the pET28a vector by GeneUniversal. Expression and purification of His-tagged UBA1 was carried out as described previously^56^.

### Steady State Kinetics Assay

Steady-state enzyme kinetic assays were performed at 30 °C in a reaction buffer containing 50 mM Bis-Tris (pH 6.5), 100 mM NaCl, and 10 mM MgClC. For the assay, varying concentrations of UBA5_37−346_ were mixed with UFM1 (5 µM) in the presence of ATP (5 µM). Additionally, UFM1 at different concentrations was combined with UBA5 (800 nM) in the presence of ATP (5 µM). Lastly, varying concentrations of ATP were tested alongside UBA5 (800 nM) and UFM1 (5 µM). The reactions were incubated with Reagent I from the AMP Glo™ kit (Promega) for 1 hour at room temperature (RT). Afterward, AMP Glo™ Reagent II was added to each well, and luminescence (Gain 150) was measured using a BioTek H1 Reader. Reaction velocities for each concentration of UBA5, UFM1, and ATP were calculated and fitted to the Michaelis-Menten equation using the GraphPad Prism 9 software. The experiment was conducted in triplicate, unless otherwise specified.

### AMP Detection Assay using the AMP-Glo^TM^ kit

The AMP Glo™ Assay Kit from Promega was utilized to detect AMP in samples following the kit’s instructions. UBA5 (800 nM) was incubated for 1 hour with various concentrations of compounds (0-100 µM). Following this, UFM1 and ATP (5 µM each) were added and incubated for an additional 30 minutes. Then, 5 µL of AMP Glo™ Reagent I was added to each well and incubated at RT for 1 hour, allowing for ATP depletion and conversion of AMP to ADP. The reaction was completed by adding 10 µL of AMP Glo™ Reagent II and incubating for 30 minutes at RT to convert ADP back to ATP, enabling the enzyme reaction. Luminescence was measured in each well using a BioTek H1 Reader, and the data were analyzed relative to a standard curve to determine AMP concentration. The ICCC values of the compounds were calculated using the GraphPad Prism 9. The experiment was performed in triplicate.

### Fluorescence Polarization (FP) Assay

To measure AMP levels in samples, the Transcreener® AMP/GMP FP Assay Kit (Bellbrook) was employed. Assay components were prepared according to the manufacturer’s instructions, with a total reaction volume of 20 µL in a 384-well black plate, consisting of 15 µL enzymatic reaction and 5 µL of kit reagent. UBA5 (1 µM) was incubated for 1 hour with varying concentrations of compounds (0-25 µM). Following this, UFM1 (2.5 µM) and ATP (20 µM) were added, and the plate was incubated for 30 minutes at RT to allow the enzymatic reaction to occur. After incubation, 5 µL of the Transcreener® AMP/GMP FP Detection Reagent was added to each well, containing the necessary fluorescent probe. The plate was incubated for 1.5 hours as specified in the protocol to develop the fluorescent signal. Fluorescence polarization was detected using a fluorescence plate reader, with the degree of polarization correlating to the amount of AMP present in the samples. The ICCC values of the compounds were calculated using GraphPad Prism 9, and the experiment was performed in triplicate.

### UBA5-free coupling Activity

Test compounds were evaluated at concentrations ranging from 0 to 100 µM. The AMP Glo^TM^ kit (Promega) was utilized for this assay with AMP as the substrate in the absence of UBA5. Following a 30-minute incubation, AMP was added to achieve a final concentration of 5 µM. Plates were read at 37 °C with a luminescence endpoint, an integration time of 2 minutes, full light emission, top optics, and gain set to 150. Appropriate controls were included. The experiment was performed in triplicate.

### UBA1 specificity assay

The UBA1 specificity assay was conducted in triplicate, following a protocol similar to the AMP Detection Assay using the AMP-Glo™ kit, which was employed for UBA5. For all UBA1 AMP-Glo assays (unless otherwise specified): UBA1, 0.3 µM; Ubiquitin, 10 µM; ATP, 5 µM.

### Charging Assay

The charging assay was conducted in duplicate as reported previously^41^. UBA5 (12 µM) was incubated with various concentrations of compounds (0-100 µM) for 1 hour in a reaction buffer comprising 50 mM Bis-Tris (pH 6.5), 100 mM NaCl, and 10 mM MgClC. UFM1 (25 µM) and ATP (2 mM) were added to the reaction samples, which were incubated for 2 hours at 30 °C. Zero time points were taken before ATP addition. Samples were analyzed *via* non-reducing 12% SDS-PAGE, followed by staining with Coomassie G-250. Product bands corresponding to each protein or complex were quantified using the ImageJ software, and percentage activity was calculated with GraphPad Prism 9.

### Inhibition of cellular UFMylation

Briefly, HEK293T cells seeded in 12 well plates (∼80% confluent) were incubated with compounds at 10 and 60 µM concentration for 24 hours, cell media were discarded, and cells were lysed using 100 µl RIPA lysis buffer. Protein concentrations were determined using BCA, western blotting experiment was performed using an anti-UFM1 antibody (Abcam, 109305; 1:2000 dilution) using 4-12% Bis-Tris Gel (Invitrogen). Quantification of the UBA5-UFM1 and UFC1-UFM1 bands was performed using ImageJ. Experiments were performed in duplicate.

### Cytotoxicity

Cytotoxicity and cell viability were measured using the cell counting kit-8 (CCK-8) (GLPBIO) according to the manufacturer’s protocol, with minor modifications. Human astrocyte cells (CCF-STTG1) were cultured in RPMI-1640 media supplemented with 10% FBS at 37 °C in a humidified atmosphere of 5% COC. A total of 1.5 × 10^5^ CCF-STTG1 cells were seeded in duplicate in a 96-well plate and incubated for 24 hours in a COC incubator. Cells were treated with UBA5 inhibitors at concentrations ranging from 0.32 to 200 μM for 48 hours at 37 °C. Following treatment, 10 μL of CCK-8 was added to each well and incubated at 37 °C for 1 to 4 hours. DMSO served as a control. Absorbance was measured at 460 nm using a BioTek Synergy HI microplate reader. The cytotoxic concentration (CCCC) was determined via nonlinear regression on dose-response curves using GraphPad Prism 9. All experiments were performed in triplicate.

## Acknowledgments

The work is supported by University of Arizona College of Pharmacy faculty startup fund, and by R. Ken and Donna Coit Endowed Chair fund in Drug Discovery. HL is additionally supported by the NIH grants: AI161845 and AI131669.

